# Ketogenic diet or BHB improves epileptiform spikes, memory, survival in Alzheimer’s model

**DOI:** 10.1101/136226

**Authors:** John C Newman, François Kroll, Scott Ulrich, Jorge J. Palop, Eric Verdin

**Affiliations:** Buck Institute for Research on Aging, Novato CA; Division of Geriatrics, University of California, San Francisco, San Francisco CA; Gladstone Institute of Virology and Immunology, San Francisco CA; Department of Chemistry, Ithaca College, Ithaca NY; Gladstone Institute of Neurological Disease, San Francisco CA

## Abstract

Links between epilepsy and Alzheimer’s disease (AD) are seen in both human patients and mouse models. Human patients with AD may commonly have subclinical epileptiform spikes (EP spikes)^1^, and overt epilepsy is associated with more rapid cognitive decline^2^. Mechanistic studies in mouse models of Alzheimer’s disease (AD) have shown that altered network activity and epileptiform spikes stem from dysfunctional inhibitory interneurons^3^, which are key elements of cortical circuits underlying cognition^4^. Treatments that reduce epileptiform spikes improve cognition in these models^5,6^. Thus, targeting subclinical epileptiform activity may be a promising new therapeutic approach to AD^7^. Ketogenic diet (KD) has long been used to treat forms of epilepsy^8^, including Dravet syndrome, a childhood epilepsy caused by mutations in a gene that is critical for inhibitory interneuron function in mouse AD models^5,9^. However, the concurrent effects of a ketogenic diet on brain electrical activity, cognitive decline, and survival have not been tested, and the translational rationale and feasibility of such an intervention remain uncertain. Here we show that a ketogenic diet reduces epileptiform spikes in the hAPPJ20 mouse model of AD. Similar reduction of EP spikes is observed using a β-hydroxybutyrate (BHB) ester in both AD and Dravet mice. A ketogenic diet improves context-dependent and visuo-spatial learning in hAPPJ20 mice. It also reduces the high seizure-related mortality observed in male mice of this model. Therapies derived from β-hydroxybutyrate may have potential application in ameliorating cognitive dysfunction in AD through reducing subclinical epileptiform activity.

---

hAPPJ20 (APP) mice carry a human APP transgene with the Swedish and Indiana familial AD-causing mutations (J20 line)^10^ and display amyloid deposition, cognitive deficits, spontaneous epileptiform spikes, and high early mortality^5,11^. In order to test if a ketogenic diet would suppress epileptiform spikes in hAPPJ20 mice, we designed a pair of diets matched for protein content and micronutrients: normal high-carbohydrate control diet based on AIN-93M (Control), and a zero-carbohydrate ketogenic diet (KD) (Fig. 1a). In pilot experiments, we found that initiation of KD did not change caloric intake (Fig. 1b), but resulted in a sustained increase of blood β-hydroxybutyrate (BHB) levels similar to that of an overnight fast (Fig. 1c). We then followed a cohort of hAPPJ20 mice undergoing serial 23-hour EEG recordings (EEGs) under three conditions: on control diet, on the second day after starting KD, and during one day of fasting (Fig. 1d). Mice recovered on the control diet for 3 weeks between the KD and fasting recordings.

**Figure 1.**
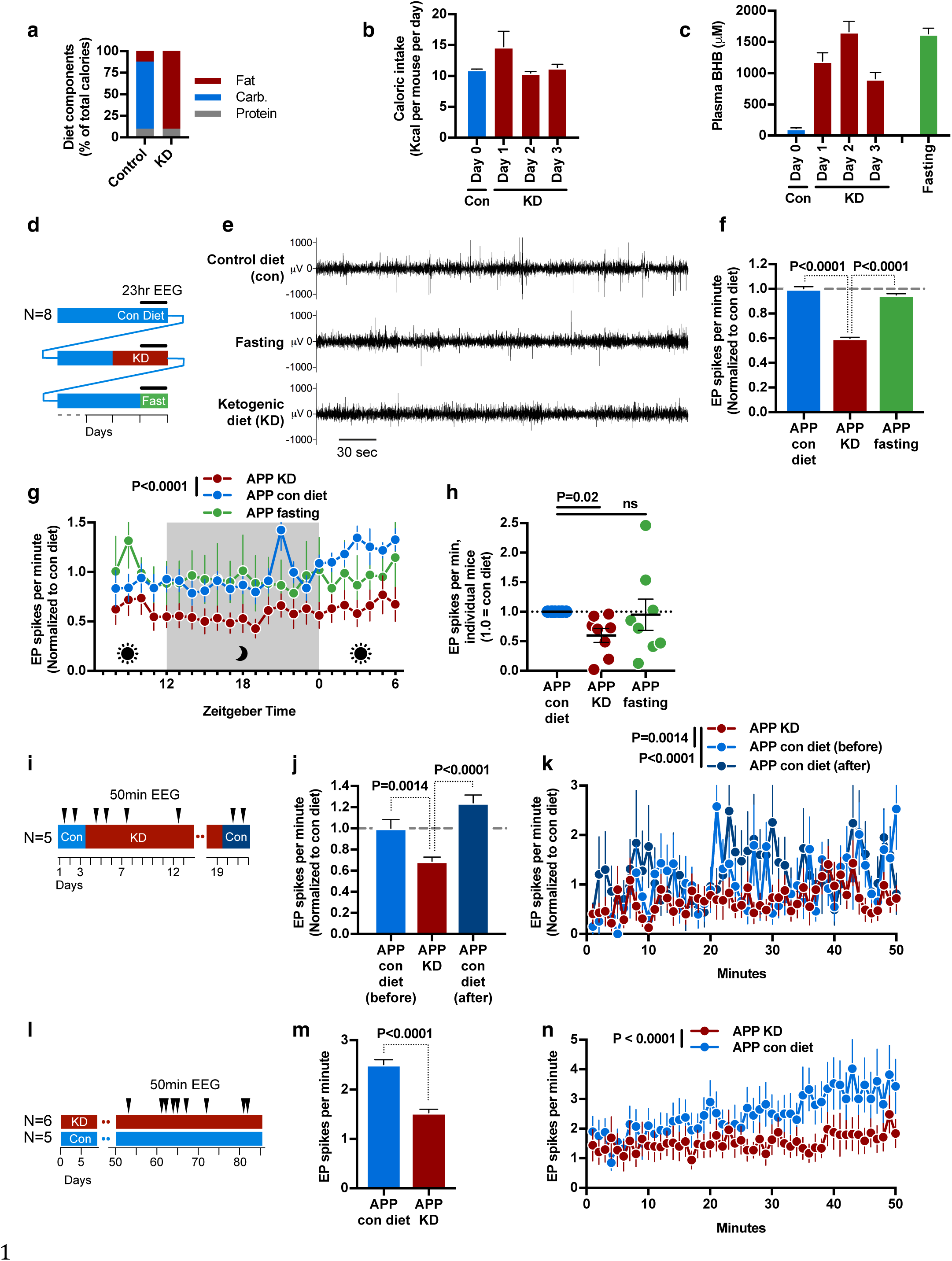
Ketogenic diet reduces epileptiform spikes (EP spikes) in the hAPPJ20 Alzheimer's mouse model. **a**, Macronutrient content of control and ketogenic diet. **b**, Initiation of ketogenic diet (KD) does not reduce caloric intake. **c**, Initiation of KD increases blood BHB levels similar to fasting. **d-h**, Longitudinal 23-hour EEG study of KD, fasting, and control diet. **d**, Study schematic. **e**, Representative EEG tracing showing EP spikes. KD but not fasting reduced EP spikes (**f**) throughout the circadian cycle (**g**). **h**, Response of individual mice to KD or fasting. **i-k**, Longitudinal cohort followed for three weeks onto and off of KD. **i**, Study schematic. KD reduced EP spikes (**j**) throughout the 50 minute recording sessions (**k**). The effect reverted within 2 days of returning to control diet. **l-n**, Separate groups of mice were followed on diets for 3 months, with 9 EEG sessions over the last 6 weeks. **j**, Study schematic. Mice on KD again had fewer EP spikes (**m,n**). In all panels, data are pooled for all EEGs within a relevant condition.

We observed that KD, but not fasting, suppressed the rate of epileptiform spikes by approximately 40% (Fig 1e and 1f). Spike suppression was consistently observed throughout the 23-hour recordings on KD (Figs. 1g). Spike suppression was also consistent between individual mice on KD, while fasting exacerbated spikes in some mice (Fig. 1h). To further explore the kinetics of EP spike suppression, we obtained EEGs with a separate cohort of mice that alternated from control diet to KD and back to control over a period of 3 weeks (Fig. 1i). Spikes were suppressed while on KD, but rebounded to prior frequency immediately after returning to the control diet (Figs. 1j and 1k). We then followed two parallel cohorts of hAPPJ20 mice fed control diet of KD for three months, with nine EEG sessions during the latter half of this period (Fig. 1l). This chronic feeding of KD suppressed EP spikes similarly to the earlier acute feedings (Figs 1m and 1n).

KD is a complex, pleiomorphic intervention with many potential mechanisms for epileptiform spike suppression, including both energetic and signaling functions of BHB, reduced insulin levels, provision of substrates for fatty acid oxidation, and many others^12^. To test if BHB is the active component of KD in spike suppression, we developed a set of novel small molecules consisting of six- or eight-carbon fatty acids ester-linked to BHB (Fig. 2a). Compound C6-BHB increased plasma BHB levels the most after intraperitoneal injection of equimolar amounts into mice fed a control diet (Fig. 2b) without altering blood glucose (Extended Data Fig. 1a). In a longitudinal, crossover study, we injected both C6-BHB and saline intraperitoneally into a cohort of hAPPJ20 mice on separate days, and recorded 50-minute EEGs both before and after each injection (Fig. 2c). Injection of C6-BHB reduced epileptiform spikes by 35% compared to saline injection (Figs. 2d) consistent with the model that BHB induced by the ketogenic diet is the key suppressor of EP spikes.

**Figure 2.**
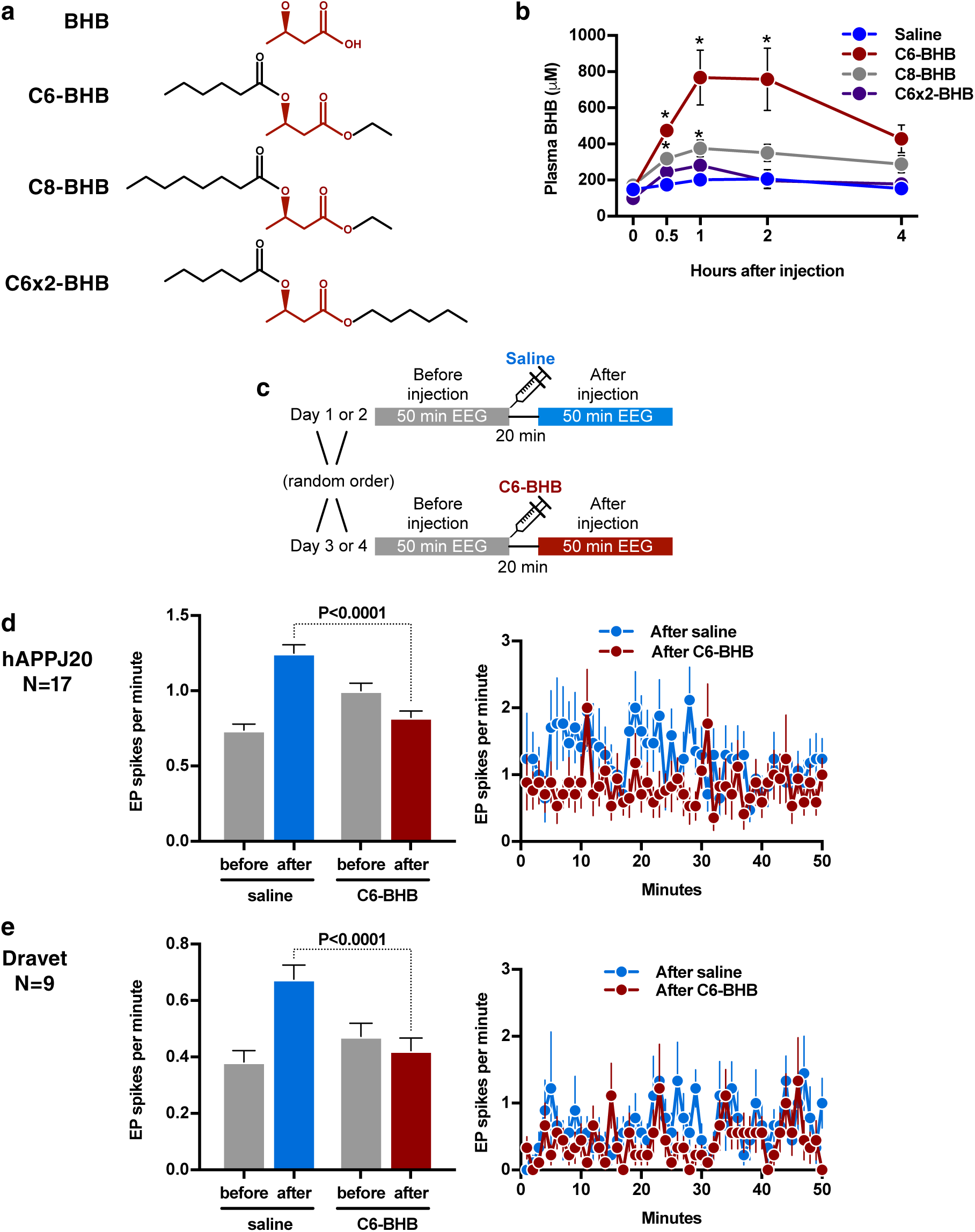
BHB ester compound reduces epileptiform spikes in hAPPJ20 and Dravet mice. **a**, Compounds with BHB ester-linked to medium chain fatty acids. **b**, C6-BHB was most effective at increasing blood BHB levels following intraperitoneal injection in wild-type mice. **c-e**, Longitudinal crossover study of C6-BHB injection **(c,** schematic) shows a reduction in spikes in both hAPPJ20 **(d)** and Dravet **(e)** mice.

Epileptiform spikes in hAPPJ20 mice is caused by the loss of activity of GABAergic interneurons which normally provide a consistent inhibitory tone and oscillatory network activity (gamma activity, measured in the 30–90 Hz gamma frequency range on EEG), thereby preventing pathological hyper synchronization of neuronal networks^13^. Interneuron dysfunction in hAPPJ20 mice in turn is associated with loss of expression of the sodium channel subunit Nav1.1 (encoded by Scn1a), and cognition is improved by restoring Nav1.1 levels by *Scn1a*-BAC expression^5^. Interestingly, Dravet syndrome, a childhood epilepsy caused by mutations in Scn1a, is often treated with a KD^9^, leading us to ask whether the C6-BHB ester would also suppress EP spikes in this mouse model. Indeed, C6-BHB suppressed EP spikes by 38% in Dravet mice (Figs. 2e) in agreement with the shared mechanistic models between Dravet and hAPPJ20 mice.

In order to assess neuronal mechanisms of spike suppression by KD, we next analyzed the relationships between exploratory movement, interneuron activity, and epileptiform spikes during the 23-hour EEG recordings (Fig. 1d). Exploratory movement is associated with increased interneuron-dependent gamma activity and suppression of epileptiform spikes in hAPPJ20^5^. However, overall movement was similar in all three conditions (Fig. 3a), indicating that spike suppression from KD was not an artifact of effects on movement. Moreover, KD reduced spikes at all movement levels, including when mice were at rest (Fig. 3b). Next we assessed whether presynaptic interneuron activity (gamma) was affected by KD. KD did not increase the induction of gamma power with increased movement (Fig. 3c), and there was no difference in overall movement-normalized gamma activity (Fig. 3d). Nor was there a change in the overall fraction of EEG power in the gamma range (Fig. 3e). Instead, we observed that mice on KD had fewer spikes at all levels of movement-adjusted gamma activity (Fig. 3f). Similar analysis of the cohort of mice that alternated from control diet to KD and back to control over a period of 3 weeks (Fig. 1i) revealed the same pattern (Extended Data Fig. 2a-c). Analysis of the BHB ester injection experiment (Fig. 2) similarly showed that the strongest reduction in EP spikes during periods at rest, when saline-injected mice had the most EP spikes (Extended Data Fig. 2d-g). Altogether, these data indicate that KD suppresses epileptiform spikes in hAPPJ20 mice; spike suppression is recapitulated by a BHB ester; and spike suppression via KD is not due to nonspecific effects on exploratory activity but occurs either independently of inhibitory interneurons, or downstream of the presynaptic potentials that generate gamma activity.

**Figure 3.**
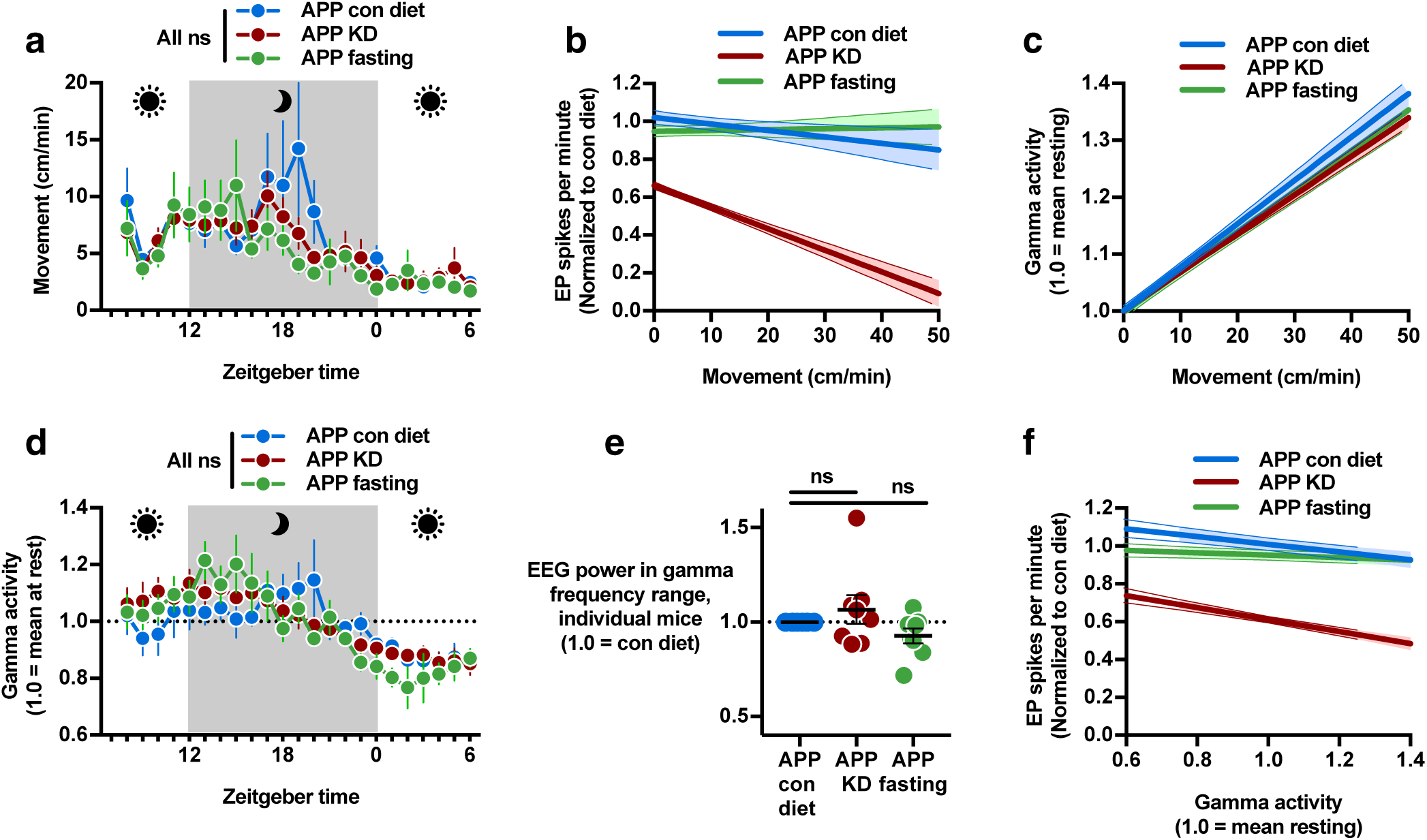
Ketogenic diet does not affect movement or interneuron gamma activity. Data were recorded during the 23 hour EEGs in Fig 1d. **a**, Movement was similar in all conditions. **b**, Regression plot of per-minute EP spikes versus movement. hAPPJ20 mice on the control diet exhibited fewer spikes during higher exploratory movement. On KD, spikes were lower at all movement levels. **c**, Regression plot of gamma activity normalized to movement (1.0 at rest) versus movement showed no change in the rate of induction of gamma activity by movement for KD or fasting. **d**, Hourly plot of movement-normalized gamma activity was not different between diet conditions. **e**, The proportion of overall EEG power in the gamma frequency range was unchanged by KD or fasting. **f**, Regression plot of EP spikes versus movement-normalized gamma activity showed fewer spikes on KD at all levels of normalized gamma.

Finally, we investigated whether EP spike suppression by KD is associated with improvements in memory or survival in hAPPJ20 mice. These mice display characteristic deficits in context-dependent learning, which can be observed as a failure to habituate to a novel environment. During a three-month study (Fig. 1l), mice were exposed to the open field four times in the first month of treatment as training, followed a month later by two test open field sessions to assess if familiarity with the open field from the prior training would reduce exploratory activity (Fig. 4a). During the test open field sessions, hAPPJ20 mice on the control diet exhibited hyperactivity and high levels of exploratory movements, as measured by total movement, center movement and rearings (Fig. 4b-d), demonstrating a lack of habituation that reflects impaired memory. Wild-type (WT) mice showed strong habituation, with low exploratory movement in the test open fields. Strikingly, hAPPJ20 mice on KD displayed habituation as strong as WT mice, with reduced total movement (Fig. 4b), and reduced exploratory movements such as movements through the center of the open field (Fig. 4c) and rearings (Fig. 4d).

**Figure 4.**
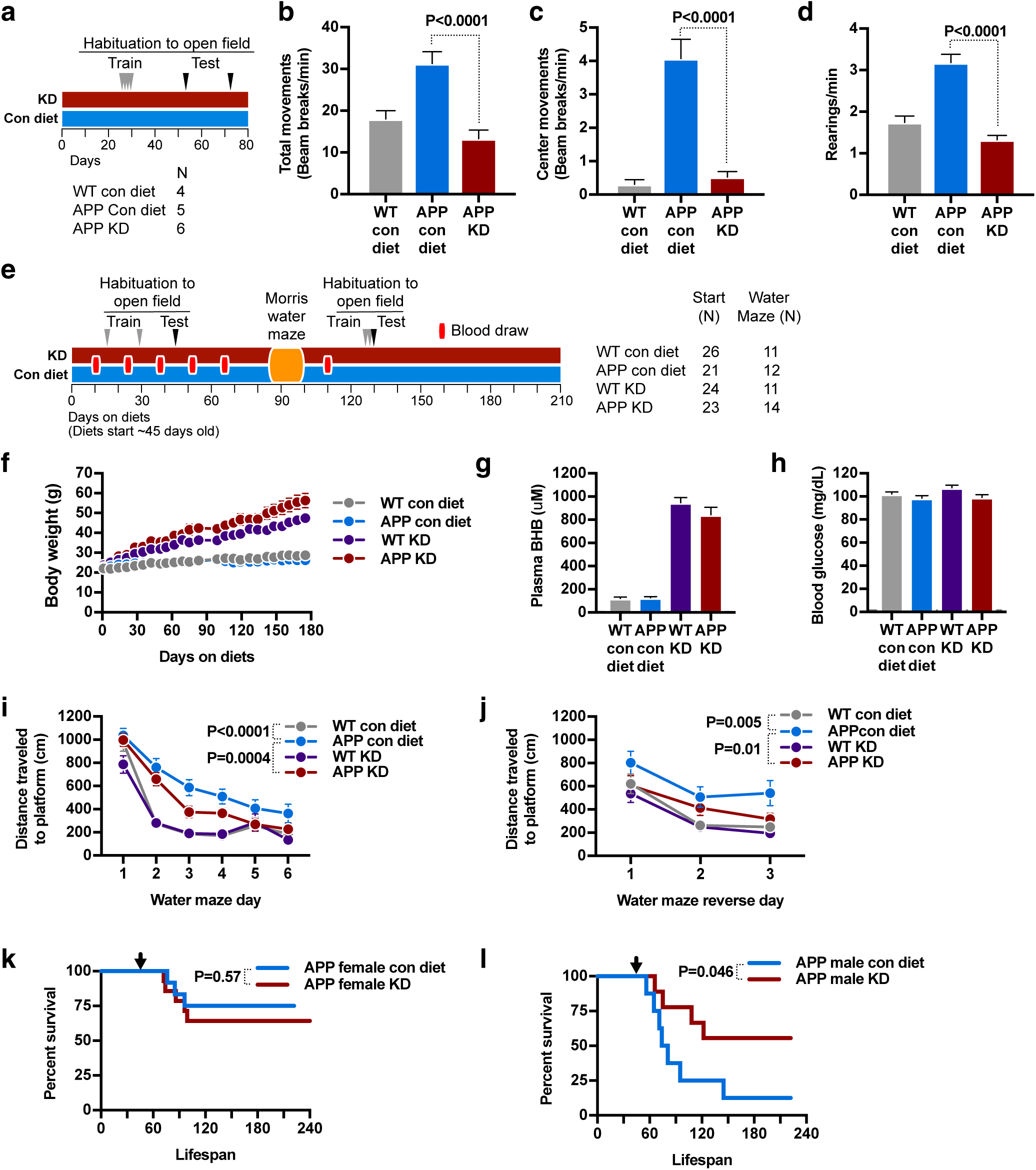
Ketogenic diet improves memory and reduces mortality in hAPPJ20 mice. **a-d**, habituation to open field context-dependent learning test on the same cohort from Fig. 1l (**a**, schematic). During the test open fields, APP mice on KD showed normal levels of movement (**b**), center movement (**c**), and rearings (**d**), indicating successful learning. Data from the two open field sessions are pooled. **e-l**, 7-month study of KD in hAPPJ20 mice **(e,** schematic). Mice gained weight on KD (**f**), showing persistently elevated blood BHB (**g**) but normal blood glucose (**h**). KD improved visuo-spatial learning performance for APP mice during both forward **(i)** and reverse **(j)** training in the Morris water maze. KD did not affect survival of female hAPPJ20 mice **(k)** but markedly improved male survival (**l**). Arrows show the start of study diets (45 days old).

To further investigate the effect of KD on memory and survival, we undertook a larger, 6-month study, beginning with 2-month-old hAPPJ20 and WT mice (Fig. 4e). KD was associated with significant weight gain in both genotypes (Fig. 4f). Six blood draws obtained every ∼2 weeks from the start of the study showed that KD was associated with plasma BHB levels averaging ∼1 mM over the 6-month period, ∼10-fold higher than controls (Fig. 4g). Blood glucose levels were similar in all groups (Fig. 4h). KD was associated with increased mean daily calorie intake over the 6 months (Extended Data Fig. 3a). Two separate studies of habituation to the open field again showed reduced exploratory center movement and rearings for hAPPJ20 mice on KD in comparison to the control diet, indicating enhanced context-dependent learning (Extended Data Fig. 3b-g). Importantly, analysis of the first open field gave no suggestion of an anxiolytic effect of KD (Extended Data Fig. 4a-d)^14^. During Morris water maze testing, hAPPJ20 mice on KD showed significantly better performance in the hidden-platform training (learning) trials of the water maze than hAPPJ20 mice on the control diet (Fig. 4i). This improvement remained consistent when the location of the platform was moved during reverse training (Fig. 4j). However, there was no difference in performance during the probe (memory) trial of the water maze, either after initial hidden platform training or after reverse training (Extended Data Fig. 5). There was no clear association between performance in the water maze and either sex or mean plasma BHB levels (Extended Data Fig. 6). hAPPJ20 mice exhibit substantial early mortality, particularly in males^5^ that may be to be due to fatal seizures. KD did not affect the relatively lower mortality among females (Fig. 4k). Remarkably, however, the high early mortality among males was strongly reduced (Fig. 4l).

In summary, we found that KD suppresses epileptiform spikes, improves context-dependent learning and visuo-spatial memory performance, and increases survival in the hAPPJ20 mouse model of AD (Extended Data Fig. 7). Spike suppression can be attained with a BHB-fatty acid ester compound on a normal diet, demonstrating that BHB is an active anti-epileptogenic component of KD in this model. These data provide a mechanistic understanding of earlier observations of visuo-spatial memory improvement with KD^15^ or a ketone ester diet^14^. The magnitude of spike suppression is similar to that obtained by low doses of the antiepileptic drug levetiracetam^6^ or by transgenic expression of SCN1A in interneurons^5^. The specific mechanism of BHB is likely multifactorial; although the immediacy of spike suppression excludes changes in Aβ accumulation, other etiologies might include mitochondrial energetics^16^, modulation of potassium channel activity^17^, deacetylase inhibition^18^, protein β-hydroxybutyrylation^19^, or increased GABA synthesis^20^. Potentiation of GABA synthesis has been observed with both BHB and KD^21^, and increased capacity for GABAergic neurons (such as the gamma-generating inhibitory interneurons) to rapidly replenish GABA from glutamate is one potential anti-epileptic mechanism of BHB that would fit our findings. While caution is appropriate in considering complex dietary interventions for elderly patients with dementia, compounds derived from BHB such as ketone esters^22^, as well as recapitulation of the molecular mechanisms of BHB, hold promise as therapies for AD. In addition, EEG monitoring of epileptiform activity could provide an important biomarker for efficacy of such agents in clinical trials.

## Acknowledgements

We thank Brett Mensh for advice and comments on the manuscript, Giovanni Maki for assistance with figure graphics, and Gary Howard for editorial assistance. Behavioral data were obtained with the help of the Gladstone Institutes’ Neurobehavioral Core (supported by NIH grant P30NS065780). This work was supported by US National Institutes of Health grants K08AG048354 (J.C.N.) and AG047313 (J.J.P.); Gladstone and Buck intramural funds (EV); funds from the Larry L. Hillblom Foundation; and Alzheimer's Association grant IIRG-13-284779 (J.J.P.).

## Author Contributions

JCN performed all experiments and data analysis except where indicated, designed the study, and wrote the manuscript. FK performed experiments in Figure 2. SU designed and synthesized compounds in Figure 2. JJP performed experiments in Figure 4, supervised experiments, and designed and supervised the study. EV designed and supervised the study, and supervised manuscript writing.

## Supplemental material

Online supplemental material includes Extended Data Figures 1-7.

## Methods

### Animal Care

All mice were maintained according to the National Institutes of Health guidelines, and all experimental protocols were approved by the University of California San Francisco (UCSF) Institutional Animal Care and Use Committee (IACUC). UCSF is accredited by the Association for Assessment and Accreditation of Laboratory Animal Care (AAALAC). Mice were maintained in a barrier facility on a 7:00 am to 7:00 pm light cycle and had free access to water. Except if stated otherwise, they were housed in littermate groups of up to 5 mice per cage, and fed ad libitum a standard chow diet (5053 PicoLab diet, Ralston Purina Company, St. Louis, MO).

### Mouse strains

hAPPJ20 mice carry a transgene containing human APP with the Swedish and Indiana FAD mutations, on a C57BL/6J background^10^. Wild-type controls are littermates that do not carry the APP transgene.

Dravet syndrome mice carry a heterozygous deletion of the Scn1a gene (Scn1a^+/-^), on a mixed C3HeB/FeJ X C57BL/6J background^23^. They were originally obtained from M. H. Meisler (Department of Human Genetics, University of Michigan), and this specific line was originally generated by K. Yamakawa (Laboratory for Neurogenetics, RIKEN Brain Science Institute). Wild-type controls are littermates that do not carry the deletion (Scn1a^+/+^).

Some ancillary experiments involving effects of ketogenic diet or novel compounds were performed using wild-type C57BL/6 male mice from the National Institute on Aging Aged Rodent Colony, usually obtained at 11 months of age, with experiments carried out between 11 and 16 months of age. These mice were tested and found to carry the Nnt partial gene deletion common to certain C57BL/6 substrains.

### Experimental diets

Customized ketogenic and control diets were obtained from Envigo. The control diet is based on AIN-93M, and contains per-calorie 10% protein, 13% fat, and 77% carbohydrates (TD.150345). The ketogenic diet contains per-calorie 10% protein and 90% fat (TD.150348). The primary fat sources are Crisco and corn oil in both diets. Fatty acids in the ketogenic diet are, by weight, approximately 24% saturated, 39% monounsaturated, and 37% polyunsaturated. The ketogenic diet is matched to the control diet on a per-calorie basis for micronutrient content, fiber, and preservatives. Note that the standard vivarium chow (PicoLab 5053) contains per-calorie 24% protein, 13% fat, and 62% carbohydrates. The ketogenic diet is of dough-like texture that permits it to be placed in the food well of the cage-top wire lid, in the same manner as pellets. All custom diets were changed weekly.

### Mouse cohort descriptions

In all figures, “APP” is hAPPJ20; “WT” are wild-type littermates that do not carry the hAPP transgene; “Con” or “con diet” is the control diet; “KD” is ketogenic diet.

Figure 1d: *N*=8 (3M, 5F), all hAPPJ20.

Figure 1i: *N=5* (2M, 3F), all hAPPJ20

Figures 1l and 4a: APP Con *n*=5 (5 M), APP KD *n* =6 (4M, 2F), WT Con *n* =4 (4 M).

Figure 2: APP *n*=17 (13M, 4F). Dravet *n*=9 (6M, 3F).

Figure 4e: At start of study, APP Con *n*=21 (9M, 12F), APP KD *n*=23 (9M, 14F), WT Con *n*=26 (12M, 14F), WT KD *n*=23 (16M, 7F). For water maze, APP Con *n*=12 (3M, 9F), APP KD *n*=14 (5M, 10F), WT Con *n*=11 (6M, 5F), WT KD *n*=11 (6M, 5F).

### Blood draws

Blood for plasma BHB testing was obtained via minimal distal tail snip, with mice placed into a whole-body restrainer (Braintree Scientific) for comfort. ∼40 μL whole blood was drawn for a BHB assay, and collected into microvettes coated with Li-Heparin (Sarstedt). Plasma was separated by centrifugation at 1500 x G for 15 min at 4 °C, and kept frozen at – 20 °C until use.

### Blood assays

BHB concentrations were determined from plasma using a BHB enzymatic detection kit (Stanbio Laboratory, Boerne, TX). Reactions were run with 3 uL of plasma each, in triplicate. Background absorbance recorded before substrate addition was subtracted form the final absorbance to correct for any hemolysis in the plasma samples. Absorbance was compared to a standard curve. We validated that a standard curve made using the kit-supplied standard (0-1 mM BHB) provided linear values up to 4 mM by testing 1-10 mM BHB solutions prepared fresh from powder (Sigma 298360). Blood glucose was evaluated on a drop of whole blood using a commercial glucometer (FreeStyle Freedom Lite).

### Electroencephalograms

Dravet and hAPPJ20 mice were implanted (under anesthesia) with Teflon-coated silver wire electrodes (0.005-inch diameter) attached to a microminiature connector bilaterally into the subdural space over frontal, central, parietal, and occipital cortices. For EEG recordings, mice were placed into clear plastic containers that were either 50 cm diameter cylinders (injection experiments) or 20x20 cm boxes (all other non-open-field recordings). Mice are free to move around the container during recordings. For 23-hour recordings, the container was lined with absorbent bedding and mice were provided with food in a small glass jar and water via gel-pack. For other recordings, the floor of the container was bare. Some EEGs were recorded during open-field experiments; this apparatus is described below. All apparatuses were disinfected with Vimoba prior to use and cleaned with 70% ethanol between mice.

EEG recording was performed with Harmonie software from Stellate Systems. Gotman spike detectors from Harmonie were used to automatically detect epileptiform spikes, which is defined by a peak lasting less than 80 ms and reaching an amplitude greater than five-fold the average baseline amplitude measured during the 5 sec before the spike^11,23^.

Mice were also videotracked during EEG recordings, and their movement quantified using Noldus Ethovision software. Videotracking during dark hours (23-hour recordings) was accomplished with infrared illuminators. Ethovision generated raw position coordinates 25-33 times per second. Despite optimization of the tracking parameters, overnight recordings in particular contained artifactual movement from centerpoint jitter and from transient aberrant detection of shadows or reflections. After extensive analysis of the movement data, we arrived at the following custom algorithm to remove artifacts: 1. Calculate a running median of raw positions across 1 second of data intervals; 2. Using the median positions, calculate the distance moved for each data interval (1/25-33 sec); 3. Ignore the distance if it fails any of three criteria: A) velocity >125cm/sec, B) direction of movement not within 90 degrees of the prior interval, C) increase in velocity >8-fold from prior interval. Although overnight (dark) recordings had the most artifacts requiring cleaning, this algorithm was applied to all movement data from all EEG experiments.

Raw power in various frequency ranges was quantified from the EEG recordings using ADInstruments LabChart software. Frequency ranges were defined as follows: 0.5-4 Hz (Delta), 4-10 Hz (Theta), 10-13 Hz (Alpha), 13-20 Hz (Beta), 30-58 Hz and 62-90 Hz (Gamma). 58-62 Hz was excluded to avoid artifacts from the 60 Hz electrical current. Power was calculated in 0.5 sec intervals, and the median of such intervals taken over each discrete minute. This per-minute power was then related to per-minute movement and spikes. The absolute magnitude of raw power is affected by various incidental factors such as length of wire, strength of wire connections, etc., and so cannot be compared directly between mice. Instead we calculated two normalized measures of Gamma frequency power: Adjusted Gamma and Gamma Fraction. Gamma Fraction is defined as the total power in the Gamma range as a fraction of the total power in all specified frequency ranges (Delta, Theta, Alpha, Beta, and Gamma). Adjusted Gamma is normalized so that 1.0 is the mean level of gamma power when the mouse is not moving. For each individual recording session (one mouse, one recording) we performed a linear regression of the set of all per-minute movement and gamma data. 1.0 is defined as the Y-intercept of the regression line, i.e. the gamma power predicted by the regression at zero mouse movement. All gamma power data points are then normalized as a simple ratio to this Y-intercept (Adjusted Gamma = [Raw Gamma] / [Regression Gamma Value at Y-Intercept]).

The several data types from the EEG experiments (epileptiform spikes, movement, and frequency power) were time-correlated for analysis using custom software.

### Open Field-EEGs and Habituation to the Open Field

The open field apparatus (automated Flex-Field/Open-Field Photobeam Activity System; San Diego Instruments) consisted of four identical clear plastic chambers (40 x 40 x 30 cm) with two 16 x 16 photobeam arrays to detect horizontal and vertical (rearing) movements. Total movements were reported, and then stratified as peripheral (4 beams on either flank of each array; 8 beams total) or central (middle 8 beams of each array). Habituation to the Open Field involves exposing mice serially to the open field environment, over which time normal/wild-type mice demonstrate reduced movement and exploratory behaviors. For EEG Open Fields, mice were prepared for EEG as above, and EEG data was collected with Harmonie software from Stellate Systems as above. Data analysis for epileptiform spikes and raw power were as above. Mice were not videotracked in the Open Field, but movement data was gathered from beam breaks.

### Morris Water Maze

The water maze pool (122 cm diameter) contained water made opaque with powdered white paint, with a 14 x 14 cm platform submerged 2 cm below the surface. For spatial training sessions, mice were trained to locate the hidden platform over at least 5 consecutive days (two sessions of two trials per day, 4 hours apart). The platform location was constant in hidden platform sessions and entry points were changed semirandomly between trials. Mice that failed to find the platform within the time limit of a training trial were placed briefly on the platform before being removed from the pool. The platform was removed for the 60 second probe trials, performed 4 hrs (followed by re-training), 24 hrs, and 48 hrs after the final training session. 24 hr probe trial data is presented. After the conclusion of the initial training and probe trials, the platform location was moved for reverse training, which involved two days of additional hidden platform training on the new location (two trials/day) followed each day 4 hrs after training by a probe trial with the platform removed. The final probe trial (after four total training trials) is presented. To control for vision and motor performance, cued training sessions with a black and white striped mast mounted above the platform were performed later. Mice were videotracked and their movement quantified during the water maze sessions with Ethovision software (Noldus).

### β-hydroxybutyrate Ester Compounds

The β-hydroxybutyrate esters were synthesized by Scott Ulrich of Ithaca College (Ithaca, New York, USA). All compounds were synthesized using the R-enantiomer of BHB as starting material, and they retain this chirality in the ester form. The common names of the compounds in Figure 4 are: hexanoyl ethyl β-hydroxybutyrate (C6-BHB), octanoyl ethyl β-hydroxybutyrate (C8-BHB), and hexanoyl hexyl β-hydroxybutyrate (C6x2-BHB). The acyl substituted ethyl β-hydroxybutyrate esters used ethyl β-hydroxybutyrate as a starting material, dissolve in pyridine and reacted with hexyl or octyl acyl chloride. The product is diluted with ethyl acetate, and washed with hydrochloric acid, sodium bicarbonate, and brine. The ethyl acetate layer is removed via drying with magnesium sulfate; the pyridine solvent is removed by rotary evaporation and vacuum pumping. ^1^H NMR and gas chromatograph mass spectrometry confirmed >95% purity of the final product. Hexyl β-hydroxybutyrate was prepared by suspending sodium β -hydroxybutyrate in dry dimethylformamide and reacting with 1-bromohexane. Washing and purification proceeded as above, except that the acid wash steps were omitted, and the purity confirmed as above before proceeding to the reaction with hexyl acyl chloride.

### Injection-EEGs

hAPPJ20 or Dravet mice were prepared for EEG as above. Mice underwent a 50 minute baseline EEG recording, followed immediately by intraperitoneal injection. They were allowed to rest 20 minutes in the home cage, then underwent a second, post-injection 50 minute EEG recording. All mice were tested with both compound and saline injections, on different days. The order of injecting compound and saline was randomized. Mice were tested in the same order on each day, so the time of day would be similar for both compound and saline injections for each mouse. Intraperitoneal injections were of 50 μL of pure compound or saline (150 mM sterile sodium chloride in water); for C6-BHB, this volume represented 0.2 millimoles of compound. Blood was drawn immediately after the post-injection EEG, or approximately 70-80 minutes after injection. Data analysis was limited to mice that completed both injections (compound and saline) and all four EEG recordings. hAPPJ20 mice were maintained on custom control diet for two weeks before and during the experiment. Dravet mice were maintained on the standard chow throughout the experiment. Note that most mice (both hAPPJ20 and Dravet) showed an increase in epileptiform spikes after saline injection, due to the stress of the procedure.

### Software

The custom software scripts used for raw data processing were written in Perl 5 with modules available on CPAN, and executed on Mac OS X. All software is available upon request and will be provided under GNU GPL licensing.

### Data analysis

Data analysis and statistical testing was performed using GraphPad Prism version 7.0. The difference between two data sets was assessed using unpaired (for most experiments) or paired (for data for which the same mouse is both control and treated at different time), two-tailed Student’s t-test. For more than two data sets, differences were assessed by one-way ANOVA with Tukey correction for multiple-hypothesis testing. Unless otherwise specified, statistical tests were performed at the level of data resolution presented on a graph - e.g. if a graph shows spikes per minute, statistical tests were performed on minute-level data. Non-significance was defined as a P-value over 0.05. Regression plots show best-fit linear regression lines with 95% confidence interval (CI) for scatterplots of per-minute data.

